# Detecting genomic deletions from high-throughput sequence data with unsupervised learning

**DOI:** 10.1101/2020.03.29.014696

**Authors:** Xin Li, Yufeng Wu

## Abstract

Structural variation (SV), which ranges from 50 bp to ∼3 Mb in size, is an important type of genetic variations. Deletion is a type of SV in which a part of a chromosome or a sequence of DNA is lost during DNA replication. Three types of signals, including discordant read-pairs, reads depth and split reads, are commonly used for SV detection from high-throughput sequence data. Many tools have been developed for detecting SVs by using one or multiple of these signals. In this paper, we develop a new method called EigenDel for detecting genomic deletions. EigenDel first takes advantage of discordant read-pairs and clipped reads to get initial deletion candidates, and then it clusters similar candidates by using unsupervised learning methods. After that, EigenDel uses a carefully designed approach for calling true deletions from each cluster. We conduct various experiments to evaluate the performance of EigenDel on low coverage sequence data. Our results show that EigenDel outperforms other major methods in terms of improving capability of balancing accuracy and sensitivity as well as reducing bias. EigenDel can be downloaded from https://github.com/lxwgcool/EigenDel.

## 1 Introduction

The differences in genetic compositions, which are relatively large in size (∼3 Mb or more) and mainly rare changes in the quantity and structure of chromosomes, are defined as microscopic structural variations [1]. With the development of molecular biology and DNA sequencing technology, smaller and more abundant alterations were observed. We define these variants, which range from ∼1 kb to 3 Mb in size, as submicroscopic structural variations [1]. Recently, they have widened to include much smaller events (for example, those >50 bp in length) [2]. The potential contribution of submicroscopic structural variants to human genetic variation and disease might be higher than that of microscopic variants, as they seem to occur at a higher frequency [1]. Deletion is a type of SVs in which a part of a chromosome is lost during DNA replication [3]. Small indels are the most common type of SVs [4]. Deletions may have significant phenotypic influence. Specifically, among genetic disorders annotated in some disease database, such as DECIPHER [5], 80% are caused by deletions [6].

Three types of alignment-based signals are used for calling deletions, including discordant read-pairs, reads depth, and split reads [2]. Many methods have been developed for SV detection by using one or multiple signals mentioned above. Pindel [7] uses an algorithm called pattern growth to report deletions with micro-insertions. Delly [8] uses split reads alignments to define the exact positions of SV breakpoints by aligning the split reads across the two regions linked by the discordant clusters, which are identified by discordant read-pairs. Lumpy [9] integrates multiple SV signals and uses different reads mappers for SV detection. Machine learning is widely used in many research fields in recent decades. Some tools, such as forestSV [10], extract the features from alignment signals and apply supervised learning method to find SV. Although many approaches have been developed for SV detection, there is no single method that outperforms others, especially in terms of balancing accuracy and sensitivity. In addition, for supervised-learning-based methods, since the benchmark repositories do not contain every SV for all individuals, the training data may contain many noises, which can significantly reduce the accuracy of prediction.

In this paper, we introduce a new unsupervised-learning-based method called EigenDel to detect germline deletions in submicroscopic SV from pair-end reads for diploid organisms. There are two major advantages of applying unsupervised-learning-based methods. First of all, since the BAM file may contain many reads mapping errors, such as repetitive ranges, it is hard to use a single threshold to separate potential deletions (homozygous/hemizygous) and normal (none-SV) ranges. Unsupervised learning can discover hidden signals within dataset, and these hidden signals are significant for calling true deletions from raw candidates. Secondly, unsupervised learning works without labeling training data, which is more adaptable than supervised learning. We compare EigenDel with other 5 widely used tools in terms of the capability of balancing accuracy and sensitivity. The results show that EigenDel outperforms these existing methods.

## 2 Method

### 2.1 High-level approach

EigenDel works with mapped sequence reads. Three statistic values, including average depth (*Depth*_*avg*_), average insert size (*Avg*_*IS*_), and standard deviation of insert size (*STD*_*IS*_) are calculated at the beginning. After that, EigenDel processes each chromosome separately to call deletions. For each chromosome, EigenDel extracts discordant read-pairs and clipped reads from mapped reads. Then, the initial deletion candidates are determined by grouping nearby discordant read-pairs. Clipped reads are used to produce more accurate estimates of the left and right breakpoints of each deletion candidate. Since the depth of deletion regions should be significantly lower than wild-type regions, candidates with depth larger than average are discarded. Then, for the remaining candidates, EigenDel gets a number of features based on depth for each of them and applies unsupervised learning to classify these candidates into four clusters. Finally, EigenDel marks these clusters as either good or bad and applies different strategies to keep true deletions from each cluster respectively. A good cluster means the majority candidates in this cluster are likely to be true deletions, while a bad cluster means the majority candidates are likely to be false. The details are illustrated in Figure 1.

**Fig. 1.**
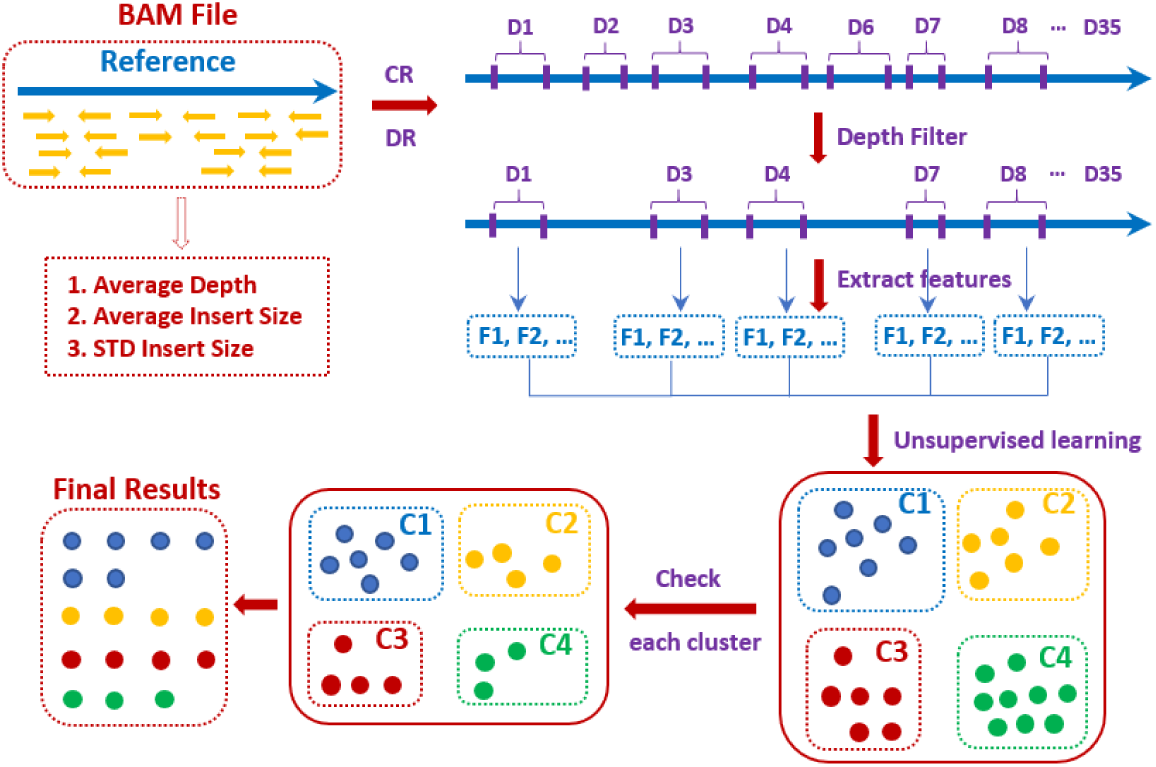
High-level approach. EigenDel takes BAM file as input. Clipped reads (CR) and discordant reads (DR) are used to obtain deletion candidates (total 35 candidates in the figure, denoted as D1 to D35). Then, some candidates, such as D2 and D6, are discarded by the depth filter. EigenDel extracts features (F1, F2, …) for each remaining deletion candidates and classify them into four clusters named C1 to C4 by unsupervised learning. There are 7, 6, 6 and 9 candidates in clusters C1 (blue), C2 (yellow), C3 (red) and C4 (green) respectively. Finally, false deletion candidates are removed from each cluster. 17 remaining candidates are called as true deletions, including 6 in C1, 4 in C2, 4 in C3 and 3 in C4.

### 2.2 EigenDel

#### Collecting border-clipped reads and discordant read-pairs

Bam file that contains alignment information of read-pairs is required by EigenDel. EigenDel uses Picard [11] to get *Avg*_*IS*_ and *STD*_*IS*_ from BAM file. Samtools [12] is used to calculate *Depth*_*avg*_. Some reads are filtered right away, including unmapped reads, polymerase chain reaction (PCR) duplicate reads, reads with low quality, and non-primary alignment reads.

Since a deletion breaks the mapping relationship between reads and reference, two types of reads, including border-clipped reads and discordant read-pairs, are collected. Border-clipped reads are the reads clipped from either tail or head, and we call them as tail-clipped reads and head-clipped reads, which are considered to support the left and right breakpoints of a deletion respectively. Since the clipped part is expected to be from the other side of a deletion, we filter the border-clipped reads, whose clipped part is shorter than 15 bp. Discordant read-pairs that satisfy *Len*_*IS*_ > *Avg*_*IS*_ + 3 ∗*STD*_*IS*_ are collected and used to locate the deletion candidates because the deletion event would enlarge the insert size of pair-end reads. Note that, since we only consider the deletion in submicroscopic structure variation, the discordant read-pairs with too large insert size are discarded. Usually, deletions do not cross different chromosomes. Therefore, we collect border-clipped reads and discordant read-pairs to identify deletion candidates for each chromosome separately.

#### Identifying deletion candidates

EigenDel first sorts all discordant read-pairs based on the position of left mates. Then it groups nearby discordant read-pairs based on the positions of their left mates to get the range of deletion candidates. Two discordant read-pairs are grouped together if the distance between their left mates is shorter than the length of read (e.g., 101 bp). Once all discordant read-pairs are grouped, each group represents a deletion candidate site. EigenDel discards candidate sites that are supported by only one discordant read-pair. The left and right boundary of each site come from the smallest mapping position of left mates and the largest position of right mates plus its alignment length respectively. Two candidate sites are merged if their boundaries are overlapped, and boundaries of the new merged site are updated. Then, EigenDel discards candidate sites that have no border-clipped reads. For each remaining site, the left breakpoint of deletion candidate comes from the largest mapping position of left mates plus its alignment length, while the right breakpoint is determined by the smallest mapping position of right mates. This roughly locates deletion candidate on the reference genome.

After that, border-clipped reads that satisfy the situations below are used to update the left and right breakpoints of deletion candidate in each site. Specifically, tail-cliped reads and head-clipped reads are viewed to contribute to left and right breakpoint respectively. For the left breakpoint, the distance between it and tail-clipped reads should be shorter than *Avg*_*IS*_. If the tail-clipped read is the second mate, its insert size should be close to *Avg*_*IS*_, and the mapping position of its first mate should be close to the left boundary of current site. If the tail-clipped read is the first mate, the mapping position of its second mate should be near the right boundary of current site. Once all qualified tail-clipped reads are collected, EigenDel only consider the best clipped positions that are supported by the largest number of tail-clipped reads. Multiple best clipped positions may be obtained, and the largest one is used to update the left breakpoint. Note we do not update it if the best clipped positions are only supported by one tail-clipped reads. There are three major differences during the updating of right breakpoint. First, the position of head-clipped reads should be near the right breakpoint. Second, if the head-clipped read is the second mate, the mapping position of its first mate should be near the left boundary of current site. If the head-clipped read is the first mate, its insert size should be around *Avg*_*IS*_, and the mapping position of its second mate should be close to the right boundary of current site. Third, the smallest best clipped positions supported by the largest number of head-clipped reads are selected to adjust the right breakpoint. Figure 2 shows the details.

**Fig. 2.**
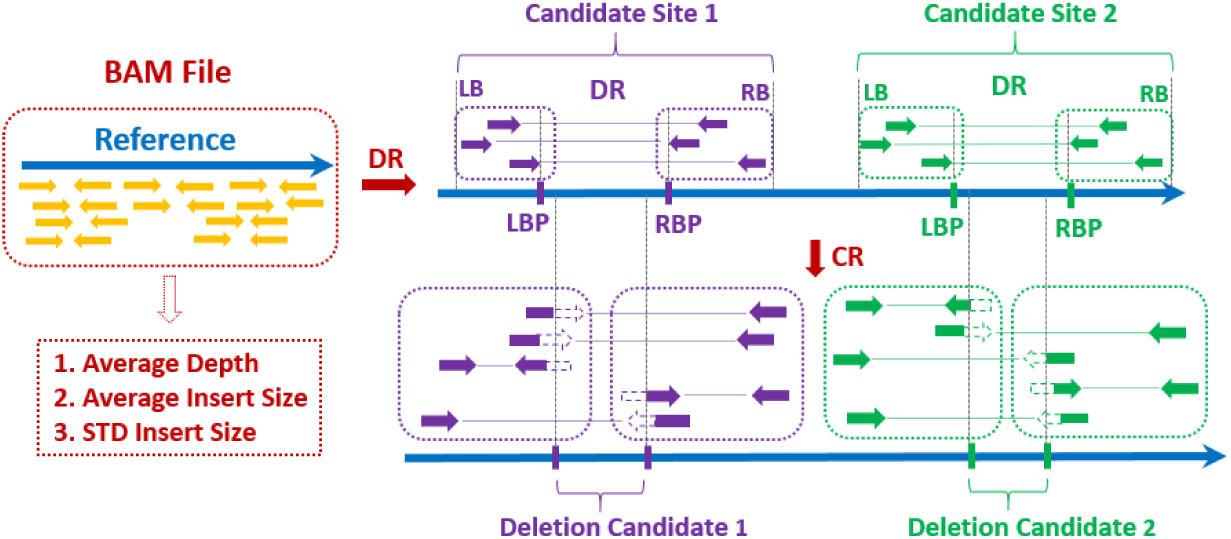
Identifying deletion candidates. Discordant read-pairs (DR) and border-clipped reads (CR) are collected from BAM file. Two deletion candidate sites, including candidate site 1 (purple) and candidate site 2 (green), are identified by DRs. Each site contains 3 DRs. Left boundary (LB) and right boundary (RB) are used to present the range of site. Left breakpoint (LBP) and right breakpoint (RBP) are used to describe the deletion candidate in current site. 5 CRs are contained by site 1, which are used to adjust LBP and RBP. Deletion Candidate 1 refers to the potential deletion in site 1. Site 2 contains 5 CRs, and its potential deletion is shown as Deletion Candidate 2.

#### Extracting features from candidates

We calculate average depth for each deletion candidate in the region between left and right breakpoints. Since a deletion may lead to significantly lower reads depth than wild-type region, the candidates with depth larger than *Depth*_*avg*_ are discarded. EigenDel is designed for detecting germline deletions in diploid organism. That is, EigenDel does not consider the situation where ploidy can change (in, e.g. tumor samples). For diploid organism, there are two types of deletions, including homozygous and hemizygous deletions. Hemizygous deletion refers to the loss of one allele, whereas homozygous (biallelic) deletion refers to the loss of both alleles identified by allele-specific analysis in clinical samples [13]. For homozygous deletions, the deletions occur in both copies. Thus, ideally, there is no reads within the deletion, and the depth should be equal to 0. For hemizygous deletion, since it is single copy deletion, the depth should be roughly equal to 50% of *Depth*_*avg*_. In practice, however, situations is less clear cut. In order to allow mapping errors and inaccurate positions of breakpoints, we identify 4 coverage ranges, namely *T*_0_, *T*_1_, *T*_2_ and *T*_3_, as shown in Table 1, to describe the internal structure of each deletion candidate.

**Table 1.** Coverage ranges for feature collection

| Coverage | Range |
| --- | --- |
| $T_0$ | 0 |
| $T_1$ | $[0, \text{Ceil}(\text{Depth}_{avg} * 0.25))$ |
| $T_2$ | $[\text{Ceil}(\text{Depth}_{avg} * 0.25), \text{Ceil}(\text{Depth}_{avg} * 0.5)]$ |
| $T_3$ | $(\text{Ceil}(\text{Depth}_{avg} * 0.5), \text{Ceil}(\text{Depth}_{avg}))]$ |

*T*_0_ refers to the perfect case of homozygous deletions (i.e., read depth is 0). *T*_1_ refers to the case of homozygous deletions allowing reads mapping errors and inaccurate boundaries. *T*_2_ refers to the case of hemizygous deletions with the same tolerance as *T*_1_. *T*_3_ refers to the range that contains both true and false deletions. We use (*D*_0_, *L*_0_), (*D*_1_, *L*_1_), (*D*_2_, *L*_2_) and (*D*_3_, *L*_3_) to present the internal structure of each deletion candidate. *L*_*i*_ stands for the total length of all positions that fall into *T*_*i*_ (may be non-consecutive), and *D*_*i*_ is the average depth of the range of *L*_*i*_. Then, we use the length of current deletion, the distance between left and right breakpoints, to normalize *L*_*i*_. We record the normalized result as *LN*_*i*_. Therefore, *LN*_*i*_(*i* = 0, 1, 2, 3) are used as 4 independent features to present each deletion candidates. Figure 3 illustrates the approach.

**Fig. 3.**
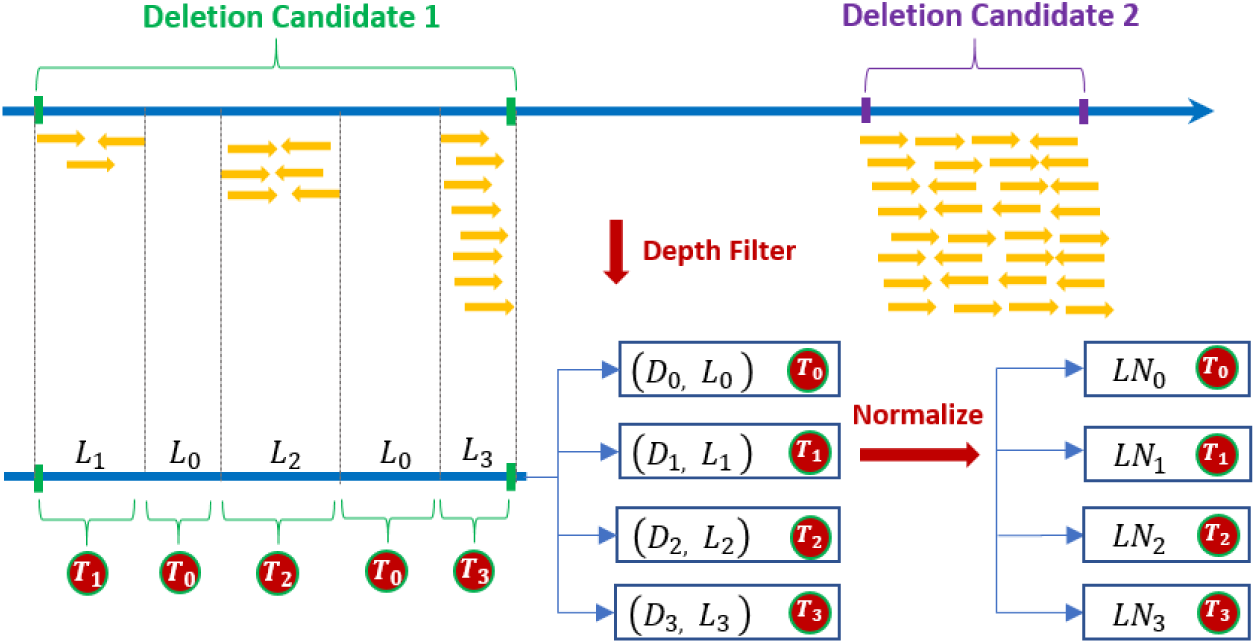
Feature extractions from deletion candidates. Two deletion candidates are identified by discordant reads. “Deletion Candidate 2” is discarded after depth filter because its depth is larger than *Depth*_*avg*_. For “Deletion Candidate 1”, 5 ranges are identified by *T*_*i*_. *L*_*i*_ and *D*_*i*_ are the total length and the average depth of the range defined by *T*_*i*_ respectively. Each *L*_*i*_ is normalized by the length of “Deletion Candidate 1”, and the normalized results are recorded by *LN*_*i*_. Therefore, the internal structure of “Deletion Candidate 1” is presented by *LN*_*i*_(*i* = 0, 1, 2, 3).

#### Detecting true deletions with unsupervised learning

So far, EigenDel collects a list of deletion candidates that are identified by discordant reads, and then the candidates are refined by clipped reads. After that, some candidates are filtered by depth filter. However there are still many false positives. For example, some false deletions may appear in the coverage range *T*_3_, which is from 50% *Depth*_*avg*_ to *Depth*_*avg*_. In addition, since the real data is noisy, it is challenging to handle some abnormal alignment situations (e.g. reads mapping error and repetitive ranges), which may change the real depth of candidates. Moreover, inaccurate breakpoints may bring the normal range into deletion candidates. These may shrink the depth difference among homozygous deletion, hemizygous deletion and normal range. Therefore, using simple thresholds alone is not be able to filter many false positives.

In order to call true deletions from noisy candidates, EigenDel applies unsupervised learning. The key idea is that different types of deletion candidates tend to cluster due to shared features. That is, the same types of true (homozygous or hemizygous) deletions tend to be similar in features (e.g., depth profile within the deletions). Similarly, same types of false positives may share some similar internal structure patterns based on reads depth. Thus, it is possible to use unsupervised learning to separate different types of deletions into different clusters. In each cluster, since the majority candidates share the similar features, it is more easy and accurate to find true deletions by applying statistical threshold. Moreover, since unsupervised learning does not need labeled samples for training, it is more flexible than supervised learning, especially for the species without good benchmark dataset.

Based on the features described in the previous step, EigenDel uses two steps to perform unsupervised learning. It first applies principle component analysis (PCA), followed by hierarchical clustering [14]. Since true deletions should be either homozygous or hemizygous, two dimensions could express all different types of true deletions. Thus, we apply PCA to all candidates and choose the top two principle components to represent each deletion. This is also good for visualization. Then, all deletion candidates are classified into four clusters based on their top two principle components through hierarchical clustering. Those clusters are expected to present 4 cases, including perfect homozygous deletions, homozygous deletions with error tolerance, hemizygous deletions with error tolerance, and the mix of heterozygous deletions and normal range. There are several advantages of hierarchical clustering. First, it does not need to select the initial node. Second, hierarchical clustering shows the relationship among the candidates in a cluster. Third, it is not sensitive to the shape of cluster (e.g. k-means prefers spherical cluster), which makes it adaptable for different dataset. The Euclidean metric and ward (sum of squares of deviations) are used for implementation.

Once four clusters are generated, they are marked as either good or bad. A good cluster means the majority of candidates in this cluster are true deletions, while the bad cluster means the majority of deletions in this cluster are false. Here is the definition of good and bad cluster. First, for a true deletion, ideally, 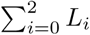 should be equal to the whole length of deletion. In another words, 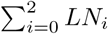 should close to 1. Considering the influence of reads mapping error and inaccurate breakpoints, we define a true deletion should have 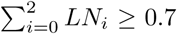. Suppose there are N deletion candidates in one cluster, we collect three values, including *LN*_0_, *LN*_1_ and *LN*_2_, for each of them. After that, all deletions in current cluster are sorted by three rounds based on *LN*_*i*_(*i* = 0, 1, 2) respectively. We record the sorted result in each round, and store them as *SR*_0_, *SR*_1_ and *SR*_2_. As a result, each *SR*_*i*_ contains all N deletions in the current cluster, which are sorted by *LN*_*i*_ from small to large. Then, we calculate three statistic values for each *SR*_*i*_, including average of 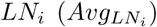, standard deviation of 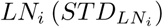 and average of top half deletions with the highest 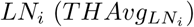. The cluster is defined as good if 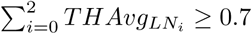, otherwise it is bad.

Once a cluster is marked as either good or bad, we use *LN*_*i*_, which is associated with the largest 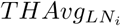, as the principle feature of current cluster to find the true deletions. We assume the distribution of *LN*_*i*_ follows empirical rule. Therefore, the majority of deletion candidates should be in the range 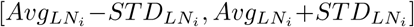, since *Pr*(*µ*−1*σ ≤ X ≤ µ*+1*σ*) ≈ 0.6827. Two thresholds, including *T*_*high*_ and *T*_*low*_, are defined by 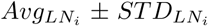 respectively. For a good cluster, the deletions are discarded if *LN*_*i*_ < *T*_*low*_ and 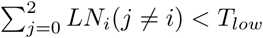. For a bad cluster, the deletions are kept if *LN*_*i*_ > *T*_*high*_ or 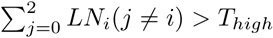. Finally, all remaining deletions in each cluster are called as true deletions. The details are shown in Figure 4.

**Fig. 4.**
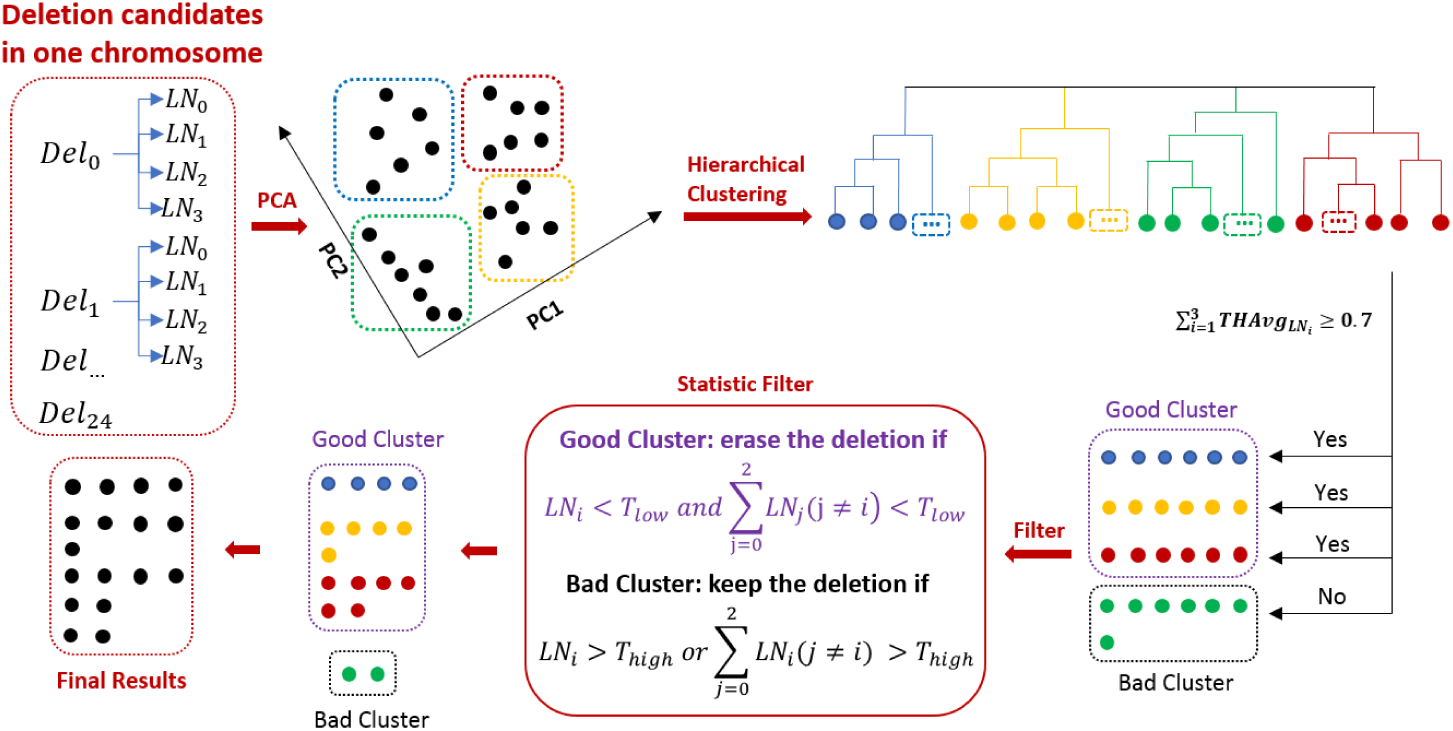
Detecting true deletions with unsupervised learning. 25 deletion candidates from *Del*_0_ to *Del*_24_ are identified, and each of which contains multiple features. PCA is applied to all candidates, and the top two principle components are used to present each candidate. All candidates are classified into four clusters through hierarchical clustering, including blue (6), yellow (6), red (6) and green (7). After checking 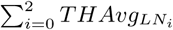, three clusters are marked as good, including blue, yellow and red, while green is marked as bad. Then statistic filter is applied to find true deletions. For a good cluster, the deletions are discarded if *LN*_*i*_ < *T*_*low*_ and 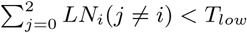. For a bad cluster, the deletions are kept if *LN*_*i*_ > *T*_*high*_ or 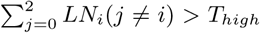. Afterwards, 4, 5, 6, 2 deletions in blue, yellow, red and green groups are remained. These deletions are reported as true deletions.

## 3 Results

We use 1000 Genome Project [15] Phase3 dataset as the benchmark, and only the deletions recorded inside are viewed as true deletions. Five existing tools are used for comparison, including Pindel, CNVnator[16], GASVpro[17], Delly and Lumpy. We directly use low coverage BAM files provided by 1000 Genome Project as input. For some tools that require separate reads files, such as Lumpy, we dump reads from BAM file.

We evaluate the performance of balancing accuracy and sensitivity by using F1 score among these methods. In our case, since there is no true negative, and all non-true positives are viewed as false positives, the precision and recall are equal to accuracy and sensitivity respectively. Therefore, the F1 score is equal to 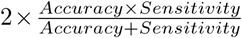 [18]. We compare F1 score based on different samples and different chromosomes in one sample respectively. A method with low bias means it can get the highest F1 score in both majority of these samples and majority of chromosomes in one sample. Our results show that EigenDel performs better than others in all testing cases.

### NA12878

The individual NA12878 in 1000 Genomes Project has been studied by many researchers. We use the low coverage BAM file (20121211) of NA12878 from the 1000 Genomes Project for comparison. The average depth of this BAM file is 5.26. It contains the aligned result of SRR622461 (92,459,459 pair-end reads). The reads length in this sequence library is 101 bps. There are 1982 deletions from 23 different chromosomes of NA12878 are reported in benchmark.

The results are illustrated in Figure 5 (A), supplemental materials (Table S1) and Figure 6 (C.1 and C.2). Figure 5 (A) shows that EigenDel has the highest F1 score for NA12878. Supplemental materials (Table S1) show that EigenDel has higher F1 score than others in the majority of chromosomes. Figure 6 (C.1 and C.2) shows an example of the performance of unsupervised learning for chromosome 1. There are 149 deletion candidates detected in chromosome 1, and 67 of them are presented in benchmark (i.e., the presumed true deletions). X and Y axes in Figure 6 (C.1 and C.2) come from the top two principle components of PCA. Figure 6 (C.1) shows all deletion candidates found by EigenDel, and the cyan dots stand for the true deletions from Phase3 callset. Figure 6 (C.2) shows the classification result of hierarchical clustering. Four clusters of deletions are generated, and they are marked in different colors. The majority of false deletions are classified in the blue cluster. The deletions in the same cluster share similar features. For example, there are 35 deletion candidates in green cluster, and the values of *LN*_0_ for all those candidates are ≤ 81%. The yellow, green and red clusters are marked as good, while the blue cluster is marked as bad. After the statistic filter is applied for each cluster respectively, 130 deletions are left (19 false deletions are discarded) and 67 of them are presented in benchmark. This means 23.2% false positives are discarded while no true deletion is lost. This demonstrates that unsupervised learning can cluster deletions with similar features, which helps to filter false positives efficiently.

**Fig. 5.**
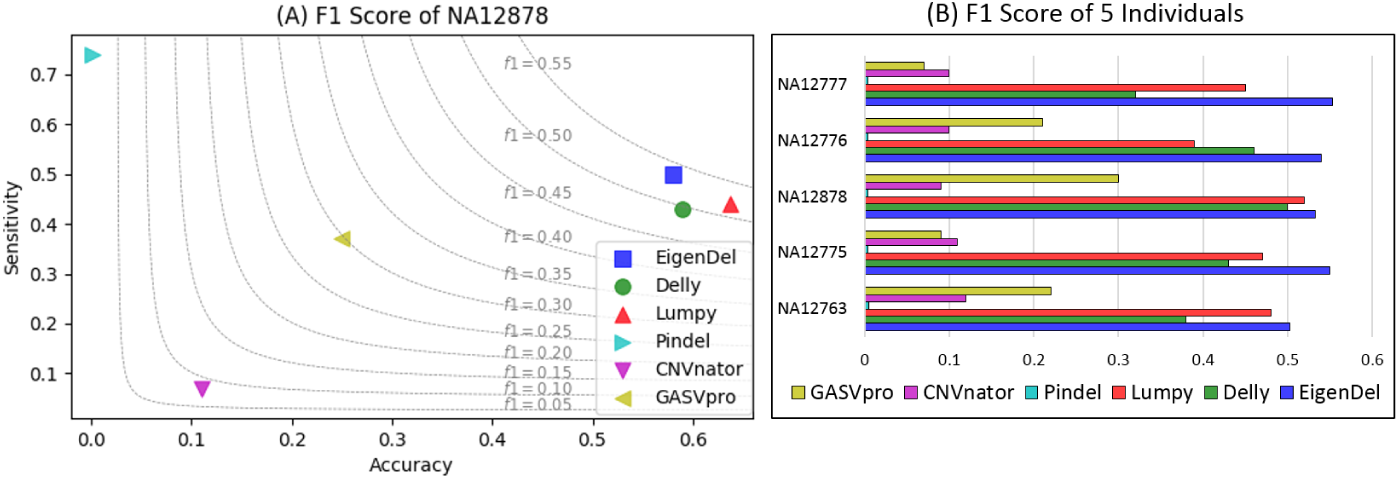
F1 scores. (A) F1 scores of all comparison tools on the whole genome of NA12878. (B) F1 scores of all comparison tools on five 1000 Genomes individuals: NA12777, NA12776, NA12878, NA12775 and NA12763.

**Fig. 6.**
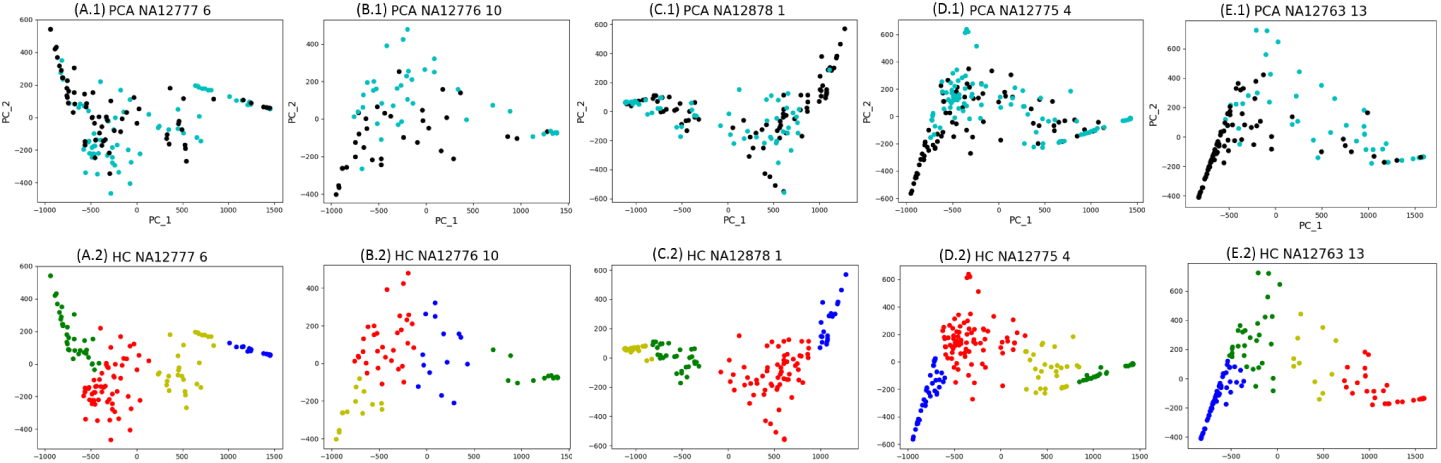
Clustering results with unsupervised learning. (A.1, B.1, C.1, D,1, E.1) The two axes are from the top two principle components of PCA. The dots represent all deletion candidates in chromosome 6, 10, 1, 4 and 13 of NA12777, NA12776, NA12878, NA12775 and NA12763 respectively. The cyan dots stand for the deletion candidates recorded in the 1000 Genomes Project Phase3 callset, which are viewed as true deletions. The black dots refer to the candidates that are not in Phase3 callset, which are viewed as false positives. (A.2, B.2, C.2, D.2, E.2) Classification results of hierarchical clustering on chromosome 6, 10, 1, 4 and 13 of NA12777, NA12776, NA12878, NA12775 and NA12763 respectively. In each scatter plot, four clusters of deletions are classified, which are marked in different colors.

### Comparison on five 1000 Genomes individuals

The low coverage BAM files from five 1000 Genomes individuals, including NA12777 (20130415), NA12776 (20130415), NA12878 (20121211), NA12775 (20130415) and NA12763 (20130502), are used in this comparison. Their reads depths are 9.08, 5.89, 5.26, 9.63 and 7.84 respectively. There are 2032, 2115, 1982, 1988 and 2105 deletions in benchmark for these five individuals respectively.

Figure 5(B) shows that EigenDel has the highest F1 score for all five individuals. Figure 6 shows the examples of clustering results of unsupervised learning from chromosomes 6, 10, 1, 4 and 13 of NA12777, NA12776, NA12878, NA12775 and NA12763 respectively. For chromosome 6 in NA12777 (Figure 6 A.1 and A.2), 140 deletion candidates are detected and 75 of them are in benchmark. After the statistic filter be applied, 23 false deletions are discarded and 71 true deletions are detected, which means EigenDel discards 35.4% false positives while only loses 5% true deletions. For chromosome 10 in NA12776 (Figure 6 B.1 and B.2), 76 deletion candidates are detected and 43 of them are recorded in benchmark. After the statistic be applied, 9 false deletions are discarded and 43 true deletions are detected, which means EigenDel discards 27.3% false positives while no true deletion is lost. For chromosome 4 in NA12775 (Figure 6 D.1 and D.2), 181 deletion candidates are detected and 103 of them are recorded in benchmark.

After the statistic filter be applied, 32 false deletions are discarded and 97 true deletions are detected, which means EigenDel discards 41% false positives while only loses 5.8% true deletions. For chromosome 13 in NA12763 (Figure 6 E.1 and E.2), 126 deletion candidates are detected and 47 of them are recorded in benchmark. After the statistic filter be applied, 50 false deletions are discarded and 46 true deletions are detected, which means EigenDel discards 63.3% false positives while only loses 2% true deletions. All results demonstrate that PCA and hieratical clustering can cluster deletions with similar features together, which helps filter false positives efficiently for different individuals on real data.

## Discussion

EigenDel is designed for detecting deletions based on low coverage data. Since low coverage data contain significant noise, it is a challenge to find true genomic deletions efficiently. We use multiple individuals, including some widely studied samples, such as NA12878, for comparison. Some BAM files contain single sequence library while others contain multiple libraries. When comparing the breakpoints of each deletion, we allow up to 15 bp tolerance.

For the performance, some tools, such as Pindel, provide high sensitivity but have a lot of false positives, which leads to low accuracy. Some other tools give better accuracy but lower sensitivity. Thus, how to balance sensitivity and accuracy is a key point of evaluation. By taking advantage of PCA and hierarchical clustering, similar deletions candidates are classified together efficiently, which helps us apply different filters to identify the true deletions in each cluster. The results show that a large number of false positives are filtered while only lose a few true deletions from the clustering results. This gives the highest F1 score among all comparison methods.

EigenDel takes 20 mins to 50 mins on running each testing sample, and this is similar to CNVnator, Delly and GASVpro. Lumpy takes around 1.5 hours on running each single individual while Pindel costs about 5 hours. As a result, the running time of EigenDel is competitive. EigneDel is designed for germline mutations of diploid organism. It uses discordant read pairs to get raw deletion candidates. Therefore, in principle, all of deletions shorter than 3 ∗*STD*_*IS*_ are discarded. The benchmark includes all types of deletions with the length from tens to tens of thousands bp. Based on the comparison results, EigenDel performs well even when short deletions in the benchmark dataset are included.

## 4 Conclusion

In this paper, we design a method named EigenDel for detecting submicroscopic structural variations deletions in germline mutation of diploid organism. EigenDel uses discordant read pairs to collect deletion candidates, and it uses clipped reads to update the boundary for each of them. The main idea of EigenDel is that it uses unsupervised learning to detect true deletions. For this, EigenDel first applies a read depth filter, and then it extracts four features for remaining candidates based on depth. Unsupervised learning is used to cluster similar deletions together: the top two principle components from PCA are used to present each deletion candidate. Hierarchical clustering is used to classify all candidates into four clusters. Then, EigenDel marks each cluster as either good or bad by using the statistic values calculated from the depth features of all candidates in the same cluster. A good cluster means the majority in the cluster are true deletions while a bad one means the majority candidates are false. EigenDel applies these different statistic filters to both good and bad clusters to extract true deletions.

The deletions from the 1000 Genomes Project Phase 3 callset are used as benchmark. The low coverage BAM files of five different 1000 Genomes individuals are used for comparison. Five existing deletion calling methods are compared with EigenDel. The results show that EigenDel gives the highest F1 score in all experiments. For each individual, EigenDel performs better than other methods in the majority of chromosomes. Thus, EigenDel has the best performance in balancing accuracy and sensitivity with low bias. EigenDel is developed by C++ and could be downloaded from https://github.com/lxwgcool/EigenDel.

## Acknowledgement

This work is supported in part by grants IIS-1526415 and CCF-1718093 from US National Science Foundation to YW.

## Supplemental materials for

### A F1 score of NA12878 in each chromosome

**Table S1.**
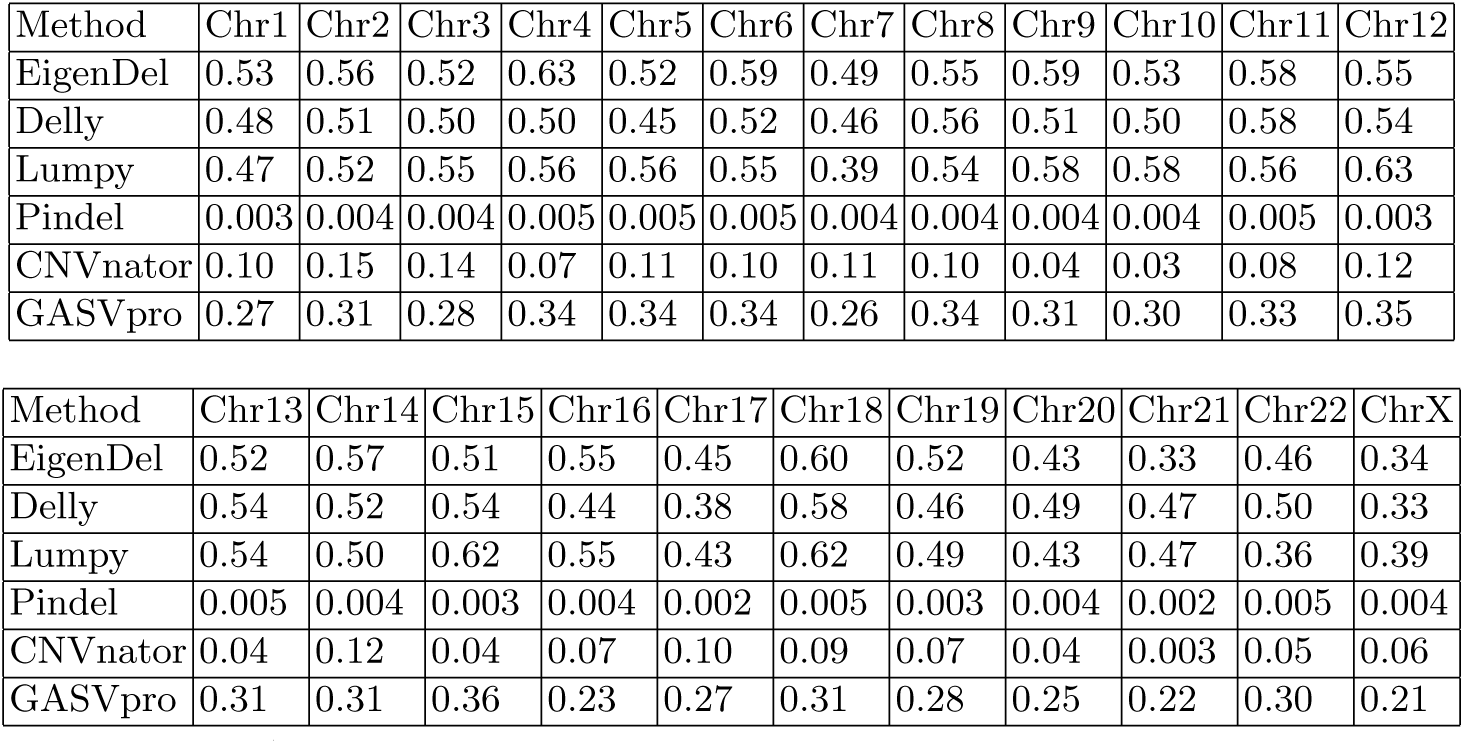
F1 score of NA12878 in each chromosome.

| Method | Chr1 | Chr2 | Chr3 | Chr4 | Chr5 | Chr6 | Chr7 | Chr8 | Chr9 | Chr10 | Chr11 | Chr12 |
| --- | --- | --- | --- | --- | --- | --- | --- | --- | --- | --- | --- | --- |
| EigenDel | 0.53 | 0.56 | 0.52 | 0.63 | 0.52 | 0.59 | 0.49 | 0.55 | 0.59 | 0.53 | 0.58 | 0.55 |
| Delly | 0.48 | 0.51 | 0.50 | 0.50 | 0.45 | 0.52 | 0.46 | 0.56 | 0.51 | 0.50 | 0.58 | 0.54 |
| Lumpy | 0.47 | 0.52 | 0.55 | 0.56 | 0.56 | 0.55 | 0.39 | 0.54 | 0.58 | 0.58 | 0.56 | 0.63 |
| Pindel | 0.003 | 0.004 | 0.004 | 0.005 | 0.005 | 0.005 | 0.004 | 0.004 | 0.004 | 0.004 | 0.005 | 0.003 |
| CNVnator | 0.10 | 0.15 | 0.14 | 0.07 | 0.11 | 0.10 | 0.11 | 0.10 | 0.04 | 0.03 | 0.08 | 0.12 |
| GASVpro | 0.27 | 0.31 | 0.28 | 0.34 | 0.34 | 0.34 | 0.26 | 0.34 | 0.31 | 0.30 | 0.33 | 0.35 |

**Table S1.** F1 score of NA12878 in each chromosome.
| Method | Chr13 | Chr14 | Chr15 | Chr16 | Chr17 | Chr18 | Chr19 | Chr20 | Chr21 | Chr22 | ChrX |
| --- | --- | --- | --- | --- | --- | --- | --- | --- | --- | --- | --- |
| EigenDel | 0.52 | 0.57 | 0.51 | 0.55 | 0.45 | 0.60 | 0.52 | 0.43 | 0.33 | 0.46 | 0.34 |
| Delly | 0.54 | 0.52 | 0.54 | 0.44 | 0.38 | 0.58 | 0.46 | 0.49 | 0.47 | 0.50 | 0.33 |
| Lumpy | 0.54 | 0.50 | 0.62 | 0.55 | 0.43 | 0.62 | 0.49 | 0.43 | 0.47 | 0.36 | 0.39 |
| Pindel | 0.005 | 0.004 | 0.003 | 0.004 | 0.002 | 0.005 | 0.003 | 0.004 | 0.002 | 0.005 | 0.004 |
| CNVnator | 0.04 | 0.12 | 0.04 | 0.07 | 0.10 | 0.09 | 0.07 | 0.04 | 0.003 | 0.05 | 0.06 |
| GASVpro | 0.31 | 0.31 | 0.36 | 0.23 | 0.27 | 0.31 | 0.28 | 0.25 | 0.22 | 0.30 | 0.21 |

### B Command lines used for genomic deletions detection

#### B.1 CNVnator

~~~
#!/bin/bash
#SBATCH -n 24
#SBATCH -N 1
#load the pre-required modules
module load gcc/5.4.0-alt zlib/1.2.11 java/1.8.0_162 mpi/openmpi/3.1.3 sam-tools/1.3 openssl/1.0.2o libcurl/7.60.0 git/2.7.2 xz/5.2.2-gcc540 bzip2/1.0.6 root/2.1
CNVnator=/scratch/xil14026/sv/software/CNVnator/0.4/cnvnator
ROOT=/scratch/xil14026/sv/data/1000Genome
BAMFILE=$ROOT/NA12763.mapped.ILLUMINA.bwa.CEU.low_coverage.20130502.bam
REFDIR=$ROOT/phase3_sv/ref_each_chrom/ref_name_std
BINSIZE=500
#Do the main logic of CNVnator
# 1: Extract read mapping
echo “Step 1: Extract read mapping”
$CNVnator -root file.root -tree $BAMFILE
# 2: Generate histogram
echo “Step 2: Generate histogram”
$CNVnator -root file.root -his $BINSIZE -chrom 1 2 3 4 5 6 7 8 9 10 11 12 13 14 15 16 17 18 19 20 21 22 X Y
# 3: Calculate statistics
echo “Step 3: Calculate statistics”
$CNVnator -root file.root -stat $BINSIZE -d $REFDIR/ # 4: Partition
echo “Step 4: Partition”
$CNVnator -root file.root -partition $BINSIZE # 5:Call CNVs
echo “Step 5: Call CNVs”
$CNVnator -root file.root -call $BINSIZE
~~~

#### B.2 Delly

~~~
#!/bin/bash #SBATCH -p general
#SBATCH -n 24
#SBATCH -N 1
module purge
module load gcc/5.4.0-alt xz/5.2.2-gcc540 bcftools/1.9 singularity/3.1
ROOT=/scratch/xil14026/sv/data/1000Genome
REF=$ROOT/phase3_sv/ref/hs37d5.fa
BamFile=$ROOT/NA12763.mapped.ILLUMINA.bwa.CEU.low_coverage.20130502.bam
DELLY=/scratch/xil14026/sv/software/delly/singularity
singularity exec $DELLY/delly_sing/ delly call -o delly.bcf -g $REF $BamFile
bcftools view ./delly.bcf > ./delly.vcf
awk ‘if($7 == “PASS”) print $0’ ./delly.vcf > ./delly_pass.vcf
~~~

#### B.3 GASVpro

~~~
#!/bin/bash
#SBATCH -n 24
#SBATCH -N 1
#SBATCH -p general
module load java/1.8.0_31 ant/1.9.4
#Main logic
GASVPATH=/scratch/xil14026/sv/software/GASV/gasv/bin
SAMPLENAME=NA12763.mapped.ILLUMINA.bwa.CEU.low_coverage.20130502
BAMFILEFOLDER=/scratch/xil14026/sv/data/1000Genome
BAMFILE=$BAMFILEFOLDER/$SAMPLENAME.bam
#1: Using BAMToGASV to Preprocess BAM files
echo “Step 1: Using BAMToGASV to Preprocess BAM files”
java -Xms512m -Xmx2048m -jar
$GASVPATH/BAMToGASV.jar $BAMFILE -LIBRARY_SEPARATED all
#2: Call structure variation
java -jar $GASVPATH/GASV.jar –batch $BAMFILEFOLDER/$SAMPLENAME.bam.gasv.in
~~~

#### B.4 Lumpy

~~~
#!/bin/bash
#SBATCH -p HaswellPriority
#SBATCH -A maa13014 #SBATCH -n 24
#SBATCH -N 1
module purge module load speedseq/0.1.2 bwa/0.7.5a python/2.7.6
module load perl/5.24.1 gcc/5.4.0-alt samtools/1.3 zlib/1.2.11 openssl/1.0.2o libcurl/7.60.0 cmake/3.8.0 python/2.7.6 pre-module post-module intelics/2017 git/2.7.2
module unload zlib
module load zlib/1.2.8-ics lumpy
SAMPLE=NA12763
REF=/scratch/xil14026/sv/data/1000Genome/phase3_sv/ref/hs37d5.fa
ROOT=/scratch/xil14026/sv/data/1000Genome/phase3_sv/Samples/$SAMPLE/reads
READ1=$ROOT/$SAMPLE.PE.1.fastq
READ2=$ROOT/$SAMPLE.PE.2.fastq
#1: generate bam file (mapped, split bam and discordant)
speedseq align -R “@RG/tID:id/tSM:$SAMPLE/tLB:lib” -t 12 $REF $READ1 $READ2
#2:run lumpy
BAM=./$SAMPLE.PE.1.fastq.splitters.bam SPLITBAM=./$SAMPLE.PE.1.fastq.splitters.bam DISCORDBAM=./$SAMPLE.PE.1.fastq.discordants.bam
lumpyexpress -B $BAM -S $SPLITBAM -D $DISCORDBAM -o sample.vcf
~~~

#### B.5 Pindel

~~~
#!/bin/bash #SBATCH -p general
#SBATCH -n 24
#SBATCH -N 1
module purge
module load samtools/1.3 zlib/1.2.11 openssl/1.0.2o libcurl/7.60.0 git/2.7.2 gcc/5.4.0-alt bwa/0.7.15 xz/5.2.2-gcc540 htslib/0.0.1 r/3.1.1 java/1.8.0_31 pdftk/2.02 bedtools/2.27.1-gcc540a
#Set variable for pindel
SAMPLE=NA12763
PINDEL=/scratch/xil14026/sv/software/pindel_new/pindel/pindel
REF=/scratch/xil14026/sv/data/1000Genome/phase3_sv/ref/hs37d5.fa
CHROM=ALL
PREFIX=./output/$SAMPLE
BAMCONFIG=./bam.config
THREAD=24
mkdir -p ./output
$PINDEL -f $REF -i $BAMCONFIG -c $CHROM -o $PREFIX -T $THREAD
~~~

#### B.6 EigenDel

~~~
#!/bin/bash #SBATCH -p general
#SBATCH -n 24
#SBATCH -N 1
module purge
module load samtools/1.3 gcc/5.4.0-alt java/1.8.0_31 sqlite/3.18.0 tcl/8.6.6.8606 python/3.6.1 r/3.1.1
../../../../MakeFile/EigenDel ./config.ini
~~~

